# Targeting the TEA DNA binding Domain of TEAD with a Peptide Inhibitor TEAi Suppresses Oncogenic Transcription and Tumor Growth

**DOI:** 10.1101/2025.08.31.672803

**Authors:** Hui Xiong, Ruohui Han, Xiaohui Lin, Shian Wu, Lei Cao

## Abstract

The Hippo pathway is a critical regulator of organ size and tumorigenesis, with the TEAD-YAP/TAZ transcriptional complex serving as a key effector frequently hyperactivated in cancers. Targeting this complex has emerged as a promising anticancer strategy. Here, we develop TEAi, a 64-amino acid peptide derived from Drosophila Nerfin-1, which directly binds the TEA DNA-binding domain of TEAD. TEAi potently suppressed TEAD-YAP-driven transcriptional activity, as evidenced by luciferase reporter and qPCR assays targeting canonical downstream genes CTGF and CYR61. Mechanistically, TEAi inhibited the DNA-binding capacity of TEAD2 without direct DNA interaction, thereby abolishing promoter recruitment of the TEAD-YAP complex. unexpectedly, TEAi induced nuclear export and cytoplasmic accumulation of TEAD2, suggesting a non-canonical mechanism distinct from native Nerfin-1 function. Functionally, TEAi significantly suppressed cancer cell proliferation in vitro and tumor growth in vivo. Unlike inhibitors targeting the YAP-binding interface, TEAi directly blocks the DNA-binding activity of TEAD, offering a potential advantage by comprehensively inhibiting all TEAD-dependent oncogenic transcription. Our findings establish TEAi as a promising prototype for therapeutics targeting TEAD, with implications for treating cancers driven by Hippo pathway dysregulation.

---

The Hippo pathway regulates organ size and tumorigenesis by balancing the homeostasis of cell proliferation and cell death. Within this pathway, the transcription co-activator YAP/TAZ lack intrinsic DNA-binding ability but interacts with DNA-binding partners such as the TEA domain (TEAD) transcription factors (TEAD1-4 in mammals) to orchestrate its transcriptional output [1-4]. Hyperactivation of the TEAD-YAP/TAZ complex is strongly linked to cancer progression, making it a promising therapeutic target [5-7].

To disrupt TEAD-YAP/TAZ signaling, we designated TEAi, a 64-amino acid peptide derived from *Drosophila* Nerfin-1 **(Figure S1A)**, which directly binds to the TEA domain of Sd, the *Drosophila* TEAD homolog [8]. To evaluate the transcriptional activity of the TEAD-YAP complex, we employed a luciferase reporter driven by the promoter of *CTGF* (Connective Tissue Growth Factor), a well-characterized target gene of TEAD-YAP in Hippo pathway signaling. Co-expression of TEAD2 and YAP in 293T cells robustly activated the CTGF-luc reporter, demonstrating functional assembly of the TEAD-YAP transcriptional complex. Strikingly, TEAi co-expression completely abolished this activation **(Figure 1A)**, indicating potent inhibition of TEAD-YAP transcriptional output. Consistence with this, quantitative real-time PCR (qPCR) analysis demonstrated that TEAi significantly suppressed YAP-induced transcriptional activation of *CTGF* as well as *CYR61*, another canonical TEAD-YAP target gene (Figure 1B and Figure S1B).

**Figure 1.**
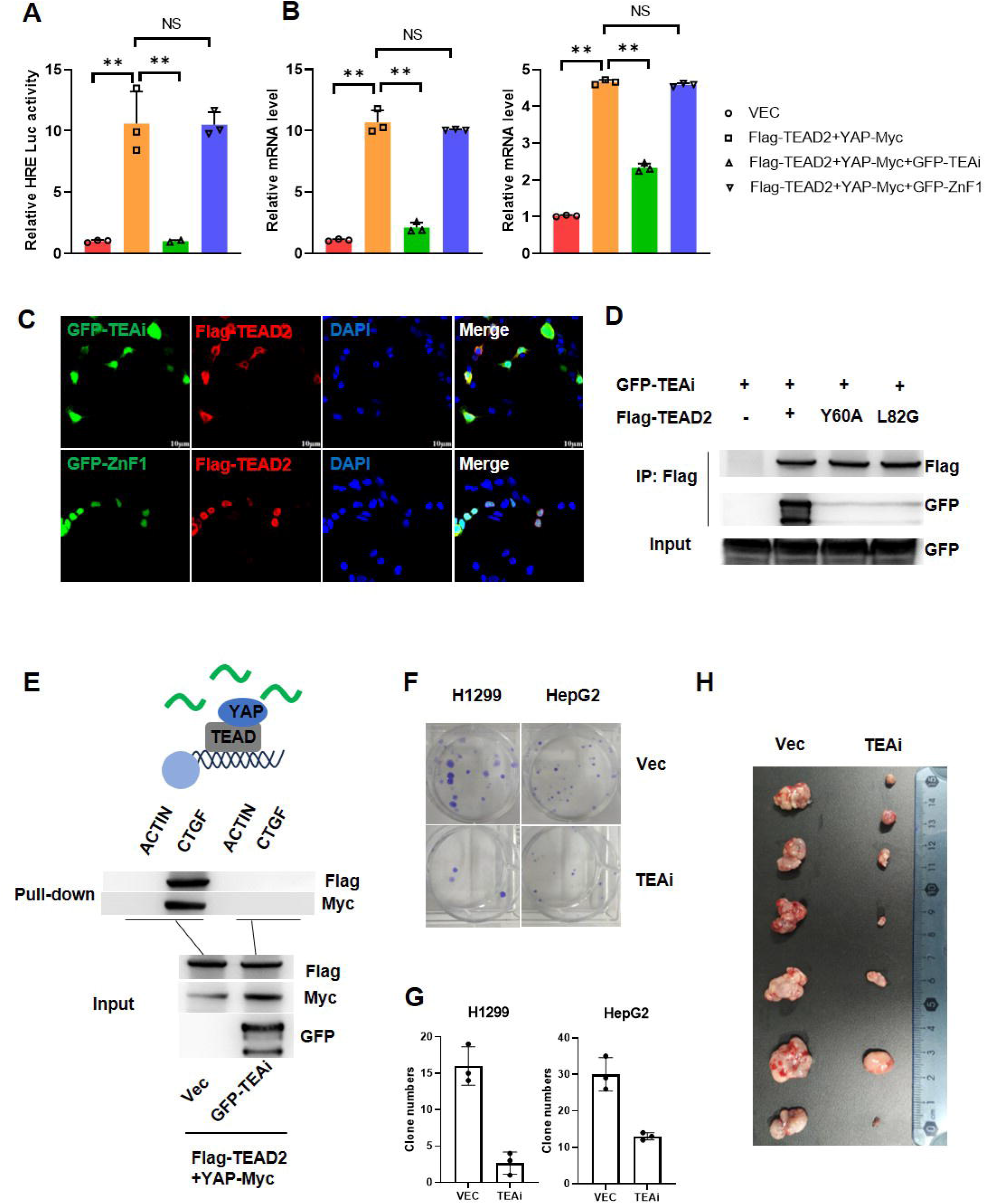
(A) 293T cells were transfected with the indicated plasmids in addition to CTGF promoter-luc plasmid and Renilla plasmid as an internal control. 36 hours later, the relative luciferase activity was measured. The error bars denote SD; n=3. P value was calculated by two-sided unpaired t test. **, P<0.01. (B) RT-qPCR analysis of the expression changes of CTGF (left) and Cyr61 (right). The error bars denote SD; n=3. P value was calculated by two-sided unpaired t test. **, P<0.01.(C) Immunostaining of 293T cells transfected with the indicated constructs. All pictures were visualized by confocal microscopy using 60’ magnification. (D) 293T cells were transient transfected by the indicated constructs and subsequently subjected to IP. WB assays to compare the levels of designated proteins in each group. (E) DNA pull-down with either ACTIN or CTGF DNA sequences from extracts of 293T cells transfected with Flag-TEAD2 and YAP-Myc with or without GFP-TEAi. WB assays to compare the levels of designated proteins in each group. (F) Colony formation assay of H1299 and HepG2 cells. (G) Quantification of (F), n=3 biological replicates. P value was calculated by two-sided unpaired t test. **, P<0.01. (H) In vivo tumor xenograft growth. Nude mice were injected with control, TEAi expressing H1299 cells and tumors were harvested after 4 weeks.

Next, we investigated whether TEAi alters TEAD subcellular localization. Surprisingly, immunofluorescence (IF) analysis revealed that TEAi promoted nuclear export of TEAD2, leading to their cytoplasmic accumulation **(Figure 1C)**. This contrasts with the canonical Nerfin-1 mechanism, which recruits CtBP corepressors to chromatin [8]. The distinct subcellular redistribution induced by TEAi suggests a novel mode of action, potentially involving disruption of TEAD-YAP nuclear retention. We thereby reasoned that TEAi may affect DNA binding of TEAD. We performed Co-IP assays in HEK293T cells expressing Flag tagged TEAD2 wild type as well as TEA domain mutants. Strikingly, TEA domain mutation abolished the interaction between TEAi and TEAD2 **(Figure 1D)**. DNA pull-down assays using a biotinylated *CTGF* promoter fragment confirmed that exdogenous TEAD and YAP specifically bound to TEAD-binding sites but not a control *ACTIN* sequence. Importantly, TEAi itself did not bind DNA but completely abolished TEAD/YAP recruitment to the CTGF promoter **(Figure 1E)**, directly linking its inhibitory effect to blockade of TEAD-DNA interaction. The DNA fragment refer to previously reported ChIP-seq data (GSM2534072, GSM1515741) to determine the TEAD binding **(Figure S1C)**.

To assess the tumor-suppressive potential of TEAi, we evaluated its effects on malignant phenotypes. Colony formation assays demonstrated that TEAi significantly inhibited cancer cell proliferation *in vitro* **(Figure 1F-1G)**. Moreover, in a subcutaneous xenograft model, TEAi-treated mice exhibited markedly reduced tumor growth compared to controls **(Figure 1H)**.

Current therapeutic strategies targeting the YAP-binding domain of TEAD (e.g., small molecules or peptides) disrupt YAP/TAZ interactions but may incompletely suppress TEAD activity, as TEAD can recruit alternative cofactors (e.g., RUNX2, Smad3, AP-1) to drive oncogenesis [9-14]. In contrast, TEAi directly inhibits TEAD’s DNA-binding ability, thereby blocking all TEAD-dependent transcriptional programs. This approach offers a potential advantage over YAP/TAZ-focused inhibitors, as TEAD knockout minimally affects normal tissue homeostasis despite its essential role in YAP/TAZ-driven tumorigenesis [1, 2, 11]. While TEAi demonstrates promising anti-tumor efficacy in vitro and in vivo, further optimization, such as enhancing peptide stability or employing nanoparticle-based delivery systems, will be critical for clinical translation.

## Methods and Materials

### Cell culture, transfection, co-immunoprecipitation, luciferase reporter assays

293T, H1299 and HepG2 cells were cultured in DMEM medium (Invitrogene) supplemented with 10% fetal bovine serum according to standard. X-tremeGENE HP DNA Transfection Reagent was used for transiently transfection according to the instructions. Transfected cells were harvested and lysed in 2×loading buffer (0.25 M Tris-HCl pH 6.8, DTT 78 mg/mL, SDS 100 mg/mL, 50% Glycerine, 5 mg/mL bromophenol blue) after 36 h for western blot. For co-immunoprecipitation, cells were lysed in NP-40 buffer (150 mM NaCl, 1% Triton X-100, 10 mM Tris pH 7.4, 1 mM EDTA pH 8.0, 1 mM EGTA pH 8.0, 0.5% NP-40, 1 mM PMSF) for 30 min at 4°C. The supernatants of the samples after centrifugation were incubated with the related antibodies for 30 min at 4°C. Samples were combined with 20 μl Protein A/G agarose (GE, 10080881) for 2 h at 4°C. Beads were washed three times with NP-40 buffer, followed by western blot. Luciferase reporter assay was performed utilizing protocols as described previously [1].

### Quantitative real-time PCR (RT-qPCR)

Total RNA was extracted using Trizol reagent (Ambion, catalog no.15596018) and subjected to reverse transcription with Superscript Reverse Transcriptase (Invitrogen, catalog no.EP0441). The RT-qPCR reactions were performed using SYBR Green Master Mix (Roche catalog no.4913914001) on 7500 Fast real-time PCR system (Applied Biosystems). The PCR conditions were performed according to the manufacturer’s instructions. The Ct values were normalized to *RPO* gene. The ΔΔCt method was used for quantifying the relative expression of target genes. RT-qPCR was repeated with three independent biological replicates.

### Immunofluorescence Staining

The immunofluorescence staining was conducted following established protocols [15] with slight modifications. Cells were first fixed in 4% formaldehyde at room temperature for 30 minutes, followed by three 10-minute washes with phosphate-buffered saline (PBS) under gentle agitation. To minimize nonspecific binding, the samples were incubated in 3% bovine serum albumin (BSA) blocking solution for 1 hour at room temperature. Primary antibodies diluted in blocking buffer were then applied to the samples and incubated overnight at 4°C. After thorough removal of unbound primary antibodies through four 10-minute PBS washes, fluorescent dye-conjugated secondary antibodies were added and incubated for 2 hours at room temperature in a light-protected environment. Following another series of four 10-minute PBS washes, cell nuclei were counterstained with DAPI (1 μg/mL) for 10 minutes and subsequently washed four times with PBS to remove excess dye. Finally, the stained samples were mounted using an antifade medium and immediately imaged under a confocal laser scanning microscope to capture high-resolution fluorescence signals.

### DNA pulldown

293T cells were seeded into 6-well plates at a density of 2 × 10□ cells per well. Each well was subjected to transfection with 500 ng of Flag-TEAD2 plasmid and 500 ng of YAP1-Myc plasmid. After 48 hours of transfection, cells were gently washed once with phosphate-buffered saline (PBS) and lysed in 500 μl of ice-cold IP buffer [50 mM Tris-HCl pH 7.5, 150 mM NaCl, 1% Triton X-100, 1 mM EGTA, supplemented immediately before use with protease/phosphatase inhibitors and 1 mM PMSF] for 10 minutes on ice. Lysates were centrifuged at 13,000 rpm for 10 minutes at 4°C to pellet cellular debris. To minimize nonspecific binding, clarified lysates were pre-incubated with 10 μg of non-biotinylated actin probe (per 100 μg of total protein) for 1 hour at 4°C under constant rotation. In parallel, 30 μl of NeutrAvidin magnetic beads (Thermo Scientific) were equilibrated with IP buffer and conjugated with biotinylated probes (*CTGF* or *ACTIN* probe) for 1 hour at 4°C with rotation. Pre-cleared lysates were then combined with the probe-bound beads and incubated overnight at 4°C with continuous rotation. Following incubation, beads were washed three times (5 minutes per wash) with IP buffer and resuspended in 30 μl of SDS loading buffer for subsequent immunoblotting analysis. DNA probes were amplified from genomic DNA using the following primers: CTGF probe (587 bp bp) F: 5’-CTCAGCGGGGAAGAGTTGTT-3’R: 5’-[Btn] ACAGGTAGGCATCTTGAGGA -3’ (5’ biotinylated) ATCB probe (558 bp): F: 5’-ctccggagatgggggaca-3’R: 5’-[Btn] aaaggcaaacactggtcgga-3’(5’ biotinylated). Short actin probe for cell extract pre-clear [(60 bp) F and R oligonucleotides were annealed at a final concentration of 1 μg/μl double-stranded DNA]:F:5’ TCGTGCGTGACATTAAGGAGAAGCTGTGCTACGTCGCCCTGGACTTCGAGCAAGAGATGG-3’R:5’CCATCTCTTGCTCGAAGTCCAGGGCGACGTAGCACAGCTTCTCCTTAATGTCACGCACGA-3’.

### Luciferase Reporter Assay

For the Dual-Luciferase reporter gene assay, the plasmids were co-transfected into 293T cells with the CTGF-Luc reporter and Renilla luciferase, which was used to normalize the transfection efficiency. The cell lysates were collected after 36 to 48 hours. The Dual-Luciferase activity was measured according to the manufacturer’s protocol (FR104-01; TransGen Biotech).

### Statistics

Statistical analysis between groups was performed by two-tailed unpaired Student’s t-test to calculate significance (*p < 0.05, **p < 0.01) for all biochemical experiments. Error bars on all graphs are presented as the S.D.

## Acknowledgments

Shian Wu and Lei Cao supervised and designed the project. Lei Cao wrote the manuscript. Hui Xiong, Ruohui Han and Xiaohui Lin performed most of the experiments. All authors read and approved the final version of the manuscript. This work was supported by the National Natural Science Foundation of China (31701126, 82300682), Natural Science Foundation of Tianjin (18JCQNJC82300) and Youth Research Incubation Fund of School of Basic Medical Sciences,Tianjin Medical University (2024FY08)..

## Conflicts of Interest

None declared.

## Notes

### Competing Interest Statement

The authors have declared no competing interest.

## Reference

1. Wu, S., et al., The TEAD/TEF family protein Scalloped mediates transcriptional output of the Hippo growth-regulatory pathway. Dev Cell, 2008. 14(3): p. 388–98.

2. Zhang, L., et al., The TEAD/TEF family of transcription factor Scalloped mediates Hippo signaling in organ size control. Dev Cell, 2008. 14(3): p. 377–87.

3. Goulev, Y., et al., SCALLOPED interacts with YORKIE, the nuclear effector of the hippo tumor-suppressor pathway in Drosophila. Curr Biol, 2008. 18(6): p. 435–41.

4. Zhao, B., Q.Y. Lei, and K.L. Guan, The Hippo-YAP pathway: new connections between regulation of organ size and cancer. Curr Opin Cell Biol, 2008. 20(6): p. 638–46.

5. Guo, P., S. Wan, and K.L. Guan, The Hippo pathway: Organ size control and beyond. Pharmacol Rev, 2025. 77(2): p. 100031.

6. Zhang, Y., et al., Mechanical forces in the tumor microenvironment: roles, pathways, and therapeutic approaches. J Transl Med, 2025. 23(1): p. 313.

7. Zhong, B., et al., The Role of Yes-Associated Protein in Inflammatory Diseases and Cancer. MedComm (2020), 2025. 6(3): p. e70128.

8. Guo, P., et al., Nerfin-1 represses transcriptional output of Hippo signaling in cell competition. Elife, 2019. 8.

9. Zhou, Z., et al., Targeting Hippo pathway by specific interruption of YAP-TEAD interaction using cyclic YAP-like peptides. FASEB J, 2015. 29(2): p. 724–32.

10. Jiao, S., et al., A peptide mimicking VGLL4 function acts as a YAP antagonist therapy against gastric cancer. Cancer Cell, 2014. 25(2): p. 166–80.

11. Liu-Chittenden, Y., et al., Genetic and pharmacological disruption of the TEAD-YAP complex suppresses the oncogenic activity of YAP. Genes Dev, 2012. 26(12): p. 1300–5.

12. Fujii, M., et al., TGF-beta synergizes with defects in the Hippo pathway to stimulate human malignant mesothelioma growth. J Exp Med, 2012. 209(3): p. 479–94.

13. Liu, X., et al., Tead and AP1 Coordinate Transcription and Motility. Cell Rep, 2016. 14(5): p. 1169–1180.

14. Chan, S.W., C. Ong, and W. Hong, The recent advances and implications in cancer therapy for the hippo pathway. Curr Opin Cell Biol, 2025. 93: p. 102476.

15. Cao, L., et al., Ubiquitin E3 ligase dSmurf is essential for Wts protein turnover and Hippo signaling. Biochem Biophys Res Commun, 2014. 454(1): p. 167–71.

