## Supplementary Figures 1 for "Targeting the TEA DNA binding Domain of TEAD with a Peptide Inhibitor TEAi Suppresses Oncogenic Transcription and Tumor Growth"

This file includes:

Supplementary Figures 1;

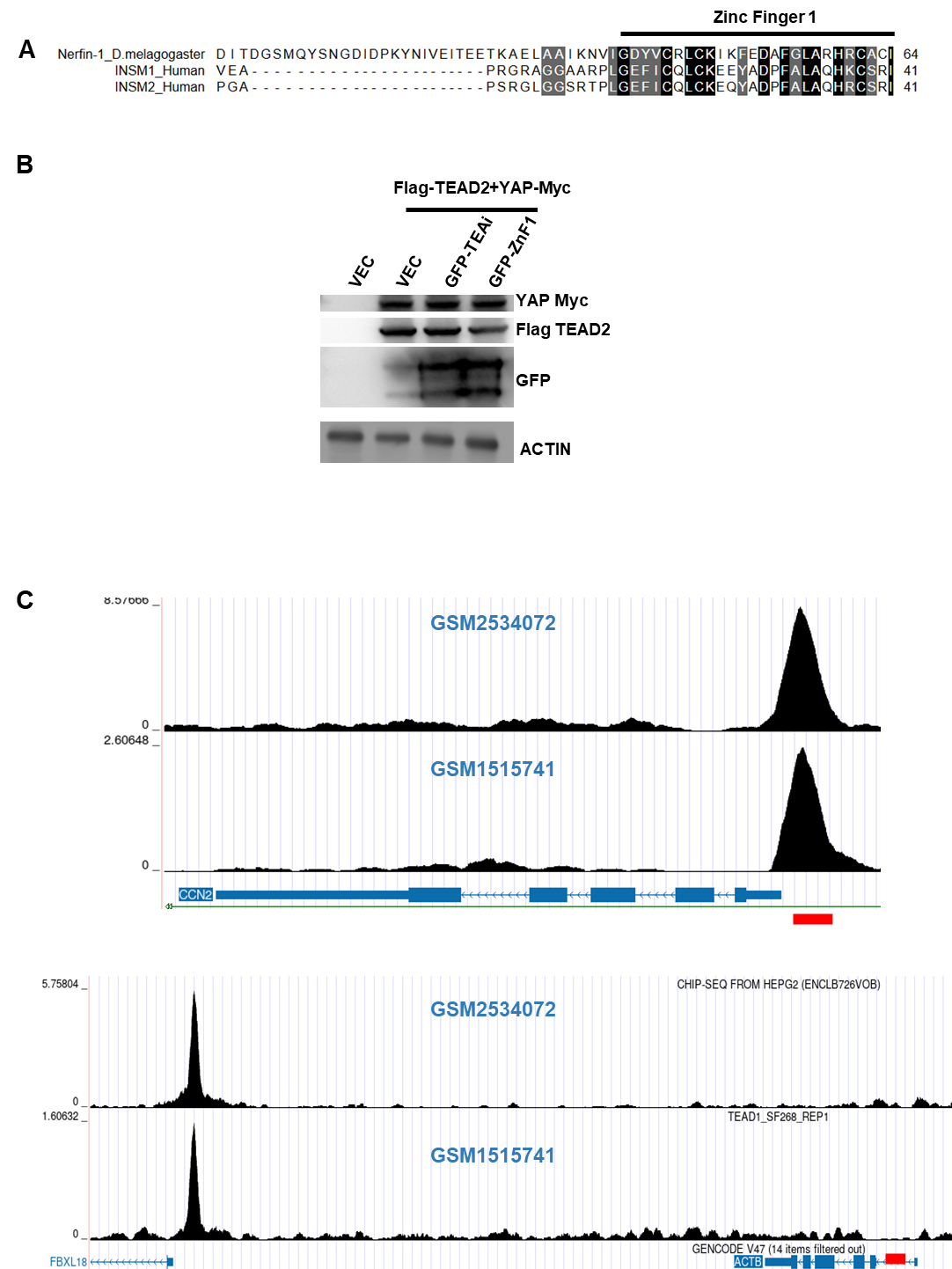
**Supplementary Figures 1**

(A) The amino acid sequence of TEAi was aligned with human homologs of Nerfin-1. (B) Western Blot assays to compare the levels of designated proteins in each group. (C) TEAD1 binding were visualized using the UCSC Genome Browser, leveraging publicly available ChIP-seq datasets (accession numbers GSM2534072 and GSM1515741) from the Gene Expression Omnibus (GEO) database.
